# Prenatal alcohol exposure programs offspring disease: Insulin resistance in adult males in a rat model of acute exposure

**DOI:** 10.1101/684639

**Authors:** Tam M T Nguyen, Sarah E Steane, Karen M Moritz, Lisa K Akison

**Affiliations:** School of Biomedical Sciences, The University of Queensland, St Lucia, QLD, Australia; Child Health Research Centre, The University of Queensland, South Brisbane, QLD, Australia

**Keywords:** fetal programming, glucose metabolism, insulin resistance, food preference, gene expression

## Abstract

Alcohol consumption is highly prevalent amongst women of reproductive age. Given that approximately 50% of pregnancies are unplanned, alcohol has the potential to affect fetal development and program chronic disease in offspring. We examined the effect of an acute but moderate prenatal alcohol exposure (PAE) on glucose metabolism, lipid levels and dietary preference in adolescent and/or adult rat offspring. Pregnant Sprague-Dawley rats received an oral gavage of ethanol (1g/kg maternal body weight, n=9 dams) or an equivalent volume of saline (control, n=8 dams) at embryonic days 13.5 and 14.5. PAE resulted in a blood alcohol concentration of 0.05-0.06% 1h post-gavage in dams. Fasting blood glucose concentration was not affected by PAE in offspring at any age, nor were blood glucose levels during a glucose tolerance test (GTT) in 6-month old offspring (*P*>0.5). However, there was evidence of insulin resistance in PAE male offspring at 6 months of age, with significantly elevated fasting plasma insulin (*P* = 0.001), a tendency for increased first phase insulin secretion during the GTT and impaired glucose clearance following an insulin challenge (*P* = 0.007). This was accompanied by modest alterations in protein kinase B (AKT) signalling in adipose tissue. PAE also resulted in reduced calorie consumption by offspring compared to controls (*P* = 0.04). These data suggest that a relatively low-level, acute PAE programs metabolic dysfunction in offspring in a sex-specific manner. These results highlight that alcohol consumption during pregnancy has the potential to affect the long-term health of offspring.

**Key points summary:**

- Prenatal alcohol exposure has the potential to affect fetal development and program chronic disease in offspring.
- Previous preclinical models typically use high, chronic doses of alcohol throughout pregnancy to examine effects on offspring, particularly on the brain and behaviour.
- In this study we use a rat model of moderate, acute, prenatal alcohol exposure to determine if this can be detrimental to maintenance of glucose homeostasis in adolescent and adult offspring.
- Although female offspring were relatively unaffected, there was evidence of insulin resistance in 6-month old male offspring exposed to prenatal alcohol, suggestive of a pre-diabetic state.
- This result suggests that even a relatively low-dose, acute exposure to alcohol during pregnancy can still program metabolic dysfunction in a sex-specific manner.

## Introduction

It is clear from both epidemiological and animal studies that early life events can impact on development and program adult disease. This is known as the developmental origins of health and disease (DOHaD) hypothesis and incorporates perturbations throughout both fetal and early postnatal development (Barker, 2007). During pregnancy, a sub-optimal uterine environment can result from maternal insults such as malnutrition, stress, obesity, and drug use. In response to a perturbation, adaptations occur in favour of vital organs, such as the brain, to ensure short-term fetal survival; however, the development of other organs deemed relatively unnecessary, such as the kidneys, liver and pancreas, can be compromised (Heindel & Vandenberg, 2015). This can result in long-term adverse health outcomes in offspring associated with the altered function in these organs. Although both clinical studies and animal models suggest that the timing of exposure can be important, organs requiring a prolonged period of development, such as the liver (Godlewski *et al*., 1997) and pancreas (Gittes, 2009), are particularly vulnerable to insults at multiple time-points. Therefore, metabolic dysfunction is commonly reported in offspring from animal models of maternal malnutrition, obesity and stress, as well as clinical studies with evidence of these exposures (see (Fleming *et al*., 2018) for review). Interestingly, effects are often sex-specific, with males consistently more susceptible to adverse outcomes than females (Weinberg *et al*., 2008; Sundrani *et al*., 2017).

One maternal perturbation receiving increasing attention is prenatal alcohol exposure (PAE). Despite health authorities advising against alcohol consumption while pregnant or planning a pregnancy (World Health Organisation, 2004; National Health and Medical Research Council, 2009), a recent systematic review and meta-analysis estimated the global rate of alcohol consumption during pregnancy to be ∼10%, although in many Western countries it is much higher (Popova *et al*., 2017). Although PAE causes well-recognised neurological and behavioural/cognitive deficits in offspring, there is also emerging evidence for a range of deficits in other body systems, leading to long-term adverse health outcomes, including impaired glucose metabolism. While clinical studies are sparse, one study reports on a small cohort (*n* = 7) of early school-age children with fetal alcohol syndrome (FAS), providing some evidence of glucose intolerance and insulin resistance in these patients compared to ‘normal’ controls (Castells *et al*., 1981). However, compelling evidence for metabolic dysfunction following PAE comes from preclinical studies. In our recent systematic review, all but three of the 18 studies included reported glucose intolerance and/or insulin resistance in PAE offspring in adulthood (see (Akison *et al*., 2019) for details). This was often associated with disruptions in molecular pathways involved in gluconeogenesis, glucose transport, IGF signalling and/or insulin signalling pathways in the liver and/or peripheral tissues. Further, recent studies, in both animal models of PAE (Amos-Kroohs *et al*., 2018; Dorey *et al*., 2018) and children with FASD (Werts *et al*., 2014; Smith *et al*., 2015; Amos-Kroohs *et al*., 2016), also report dysregulated eating behaviours and food preference, potentially contributing to a propensity towards diabetes or other metabolic dysfunction and associated comorbidities such as obesity.

However, in most of these preclinical models, chronic high doses of ethanol were administered, resulting in daily peak blood alcohol concentrations (BAC) of 100-150 mg/dl (0.10-0.15%) (Chen & Nyomba, 2003b). This is not representative of drinking patterns reported in the majority of pregnant women, which are typically acute, low-moderate levels of alcohol, referred to as ‘special occasion’ drinking (Muggli *et al*., 2016; McCormack *et al*., 2017). Therefore, although the effects of substantial ethanol consumption are well-recognised, the effects of mild exposure are less well known.

In the current study, we used a rat model of moderate, acute alcohol exposure during an important stage for liver and pancreas development, and when fetal development is equivalent to that of the first trimester human fetus, when the mother may still be unaware of her pregnancy. Timing of exposure was based on the relative development of these organs between humans and rodents (Godlewski *et al*., 1997; Pan & Brissova, 2014). Food preference for a HFD compared to standard chow was examined at 4-5 months of age and then a glucose tolerance test (GTT) and insulin tolerance test (ITT) were performed in offspring at 5-6 months of age. In order to elucidate potential molecular mechanisms contributing to disease phenotypes, the effect of prenatal alcohol exposure on hepatic expression of genes involved in glucose homeostasis and AKT-signalling in peripheral tissues were also examined in adult offspring. We hypothesised that our moderate, acute prenatal alcohol exposure could program glucose intolerance and insulin resistance in adult offspring, as well as a preference for a high-fat diet. Given previous reports of sex-specific effects, we included sex as a potential contributing factor in all analyses.

## Materials and Methods

### Ethical approval

All animal experiments and procedures were approved by The University of Queensland Anatomical Biosciences Ethics Committee (SBMS/AIBN/521/15/NHMRC) and were conducted in accordance with the Australian Code for the Care and Use of Animals for Scientific Purposes (2013, 8th Edition). Reporting of animal experiments conforms to the ARRIVE guidelines (Kilkenny & Altman, 2010; Kilkenny *et al*., 2010), as well as the principles and regulations for animal experiment reporting and ethics described by (Grundy, 2015).

### Animal model of acute prenatal alcohol exposure

Outbred, nulliparous Sprague-Dawley rats were obtained from the Animal Resources Centre, Perth WA. All animals (F0 and F1) were housed at the Australian Institute for Bioengineering and Nanotechnology (AIBN) Animal Facility (University of Queensland, St Lucia, QLD, Australia) in a temperature- and humidity-controlled environment with an artificial 12 h reversed light-dark cycle and provided with standard laboratory rat chow (Rat & Mouse Meat-Free Diet, Specialty Feeds, Glen Forrest WA, Australia) and water ad libitum. Following transport, females were group-housed during an initial acclimation period in open-top cages with wire lids, plastic bases and wood-chip bedding.

At a weight threshold of 230 g (∼3-4 months old), dams were assessed daily for proestrous via vaginal electrical impedance (≥4.0 kΩ) using an EC40 estrous cycle monitor (Fine Science Tools, Foster City, CA, USA) as previously described (Jaramillo *et al*., 2012). Once in proestrous, dams were housed with a proven stud male for the first 5 h of the dark cycle (1200 – 1700) and successful mating confirmed by the presence of a seminal plug, with the following morning designated as embryonic day (E) 0.5. Pregnant dams were housed singly. Dams were weight-matched and then randomly assigned to either receive ethanol (EtOH) or saline (Control) via oral gavage at E13.5 and E14.5. EtOH treated females (*n* = 10) received 18% v/v EtOH in saline solution (0.9% NaCl) at a dose of 1 g/kg body weight, while Control females (*n* = 8) received an equivalent volume of saline. Gavage was performed between 09:00-10:00 each day (i.e. during the light cycle). Water and chow consumption were measured daily from E12.5 (one day prior to first gavage) until birth. Weight gain was monitored throughout pregnancy. Day of birth was designated postnatal day (PN) 0, with offspring weighed on alternate days from PN1, then weekly following weaning at PN21. Once offspring were weaned, all dams were culled via CO_2_ asphyxiation. Only 1 male and 1 female were used from each litter for each experiment to remove potential litter effects, as recommended in recent guidelines on experimental design in DOHaD studies (Dickinson *et al*., 2016). Where >2 pup weights were measured per litter, litter averages were calculated and used in subsequent analyses. Figure 1 provides a flow chart showing the dams allocated to each arm of the study and offspring that contribute to each analysis at each age. All procedures conducted on offspring were performed during the light cycle (0930-1200).

**Figure 1:**
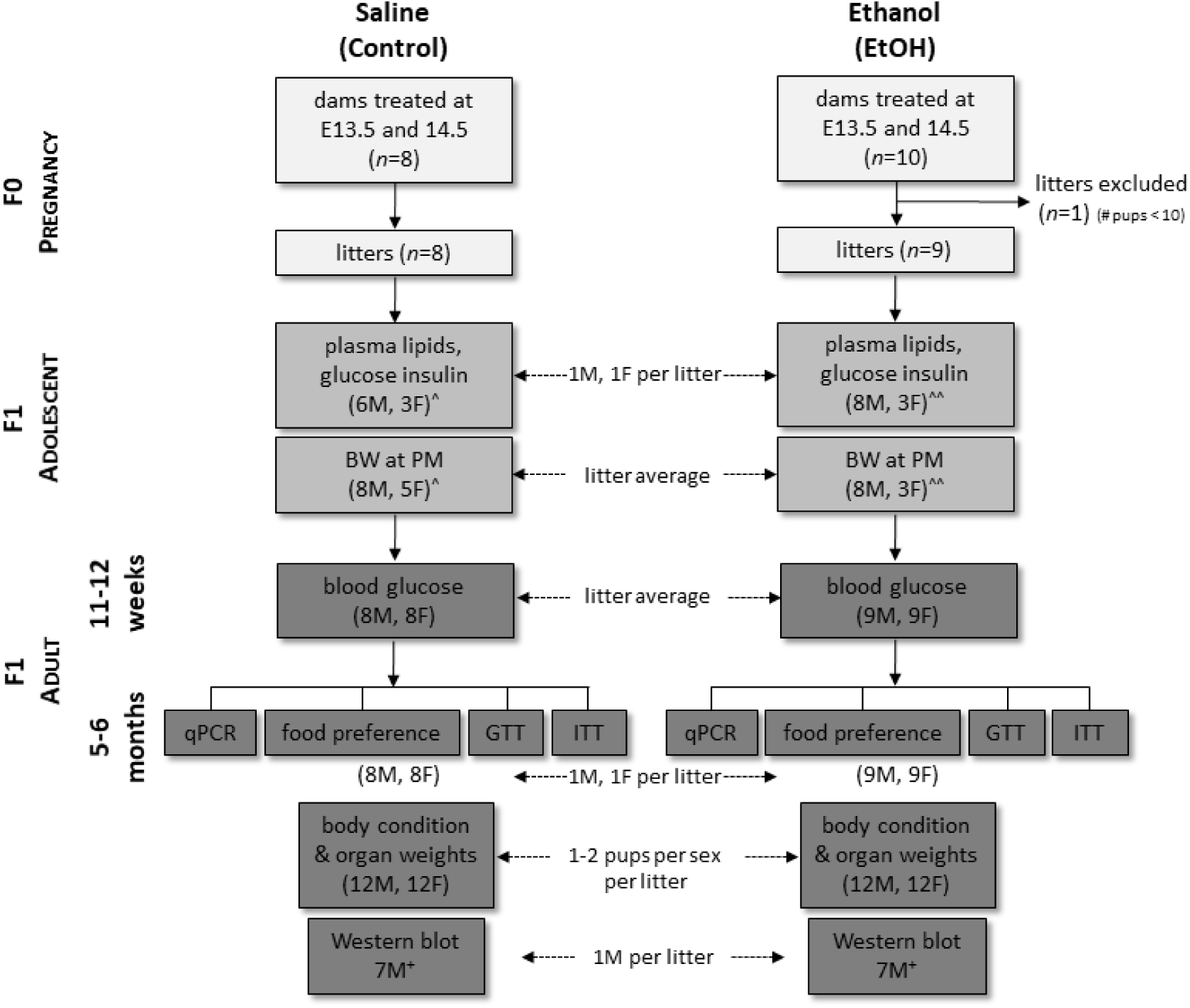
Flow chart of animals treated and offspring used in each experiment. Pregnant dams were treated with either saline or 1g/kg body weight ethanol (EtOH) via gavage. Note that one EtOH litter was excluded from further experiments given the litter size was 7 (mean, range for other litters: 15, 11-19). ^^^ <8 females available due to litter size/sex-ratio constraints and use in another study; available females were prioritised for adult experiments; 2 litters could not be used in plasma analyses as not fasted. ^^^^ <9 females available due to litter size/sex-ratio constraints and use in another study; available females were prioritised for adult experiments; 1 litter had no animals used at this time-point. ^+^ Only males analysed; sample size reduced to allow all samples to be run on one gel. F = female; M = male; E = embryonic day; BW = body weight; PM = post-mortem; GTT = glucose tolerance test; ITT = insulin tolerance test; qPCR = real-time quantitative polymerase chain reaction.

### Blood and tissue collection

Approximately 150 μl whole blood was collected from dams via a tail tip bleed at 1 h and 5 h post gavage on E13.5 and 14.5 to measure blood alcohol concentration (BAC; see details for measurement below) and non-fasted blood glucose level using an Accu-Chek Performa glucometer (Roche, Bella Vista, NSW, Australia). At PN30, a subset of offspring (see Figure 1) were weighed and then culled via CO_2_ asphyxiation following a 16 h fasting period. Blood was collected via cardiac puncture for measurement of fasting plasma insulin, metabolites involved in lipid metabolism and fasting blood glucose. Blood was immediately centrifuged at 4000 rpm for 10 min at 4°C and plasma separated, aliquoted and stored at -20°C until subsequent analysis. Fasting blood glucose was also measured, following a 16 h fasting period, in all offspring at 10-11 weeks of age via a spot tail bleed and glucometer. Litter-mate blood glucose levels were averaged for each sex prior to analysis for differences between PAE and control. At 6 months of age, 1 male and 1 female per litter had blood collected as part of a glucose tolerance test (GTT) or insulin tolerance test (ITT) (see below for details). All offspring were culled at 6-7 months of age via CO_2_ asphyxiation, with a subset used for tissue collection (1-2 males and 1-2 females per litter for a total of 12 per sex per treatment). Females were assessed for proestrous (≥4.0 kΩ vaginal impedance) as described above and were not culled at this stage of the cycle to reduce potential variability due to hormonal changes just prior to ovulation. Body weight, abdominal circumference, snout-rump length and tibia length were measured for calculations of size and ponderal index (weight in g/cubed length in cm). Pancreas and liver were removed, weighed and a section from the largest lobe of the liver snap-frozen in liquid N_2_ and stored at -80°C for subsequent molecular analysis. Samples of abdominal visceral fat and the left gastrocnemius muscle were also frozen for subsequent molecular analysis.

### Measurement of blood alcohol concentration (BAC)

Maternal blood samples were immediately centrifuged after collection at 4000 rpm for 10 min at 4°C and plasma separated, aliquoted and stored at -20°C until analysed. Blood alcohol concentration (BAC) was measured by Pathology Queensland (Queensland Health) using an alcohol dehydrogenase enzymatic assay (Beckman Coulter, Ref #474947) and a Beckman Coulter AU480 Chemistry Analyzer (Beckman Coulter, Lane Cove, NSW, Australia). The limit of detection for this assay was 5 mg/dL or 0.005%.

### Food preference study

Food preference was assessed in offspring 4-5 months of age as previously reported (Dorey *et al*., 2018). Briefly, 1 male and 1 female from each litter were randomly chosen and housed individually in cages with a divided feed hopper. After a 4-day acclimatisation period, baseline consumption was established by measuring intake of standard chow diet (SD, 4.8% fat) daily for a period of 4 days and calculating consumption per gram of body weight. Offspring were then allowed free access to both a high-fat ‘Western’ style chow diet (HFD; 21% fat; Specialty Feeds Diet SF00-219) and the SD over a 4-day test period. The diets were randomly placed on either side of the feed hopper and their positions switched after 2 days. Cage positions were also randomised on the racks and were changed after 2 days to prevent systematic sampling bias of Control and EtOH animals. Food and water intake were recorded daily and consumption per gram of body weight calculated over the first 2 days (Choice Period 1, CP1) and the second 2 days (Choice Period 2, CP2) of the two diets being offered. The test period was split into CP1 and CP2, as a previous study has shown that the novelty of the HFD diminishes over the test period, resulting in different eating behaviour over the 4-day period (Dorey *et al*., 2018).

### Glucose and insulin tolerance tests

At 6-7 months of age, a GTT and ITT were performed as previously described (Probyn *et al*., 2013; Gardebjer *et al*., 2015). Briefly, 1 male and 1 female were randomly selected from each litter (EtOH = 9; Control = 8) for each test. For the GTT, offspring were fasted overnight for 12-16 h and then received an intraperitoneal (ip) injection of a 50% w/v glucose solution (Baxter Healthcare, Old Toongabbie, NSW, Australia) at a dose of 1 g/kg body weight. Blood was sampled via tail tip bleed, and blood glucose concentrations were measured using a glucometer prior to glucose administration (-5 min), and at 5, 10, 20, 30, 45, 60 and 90 min post-bolus. Blood was also collected, processed as described above and plasma stored for subsequent analysis of plasma insulin and glucose levels. For the ITT, a separate group of non-fasted offspring received an ip injection of 0.75 U/kg body weight insulin (Actrapid, Novo Nordisk Pharmaceuticals Pty. Ltd., Baulkham Hills, NSW, Australia), and tail tip bleeds were performed to measure blood glucose concentrations via glucometer as above at -5, 20, 40, 60, 90 and 120 min post-injection.

### Quantitative PCR and Western blotting

Expression of genes involved in hepatic glucose metabolism and peripheral tissue glucose transport and insulin signalling were examined using real-time quantitative polymerase chain reaction (qPCR) (see Table 1 for details of specific genes). qPCR was conducted and reported following the MIQE guidelines (Bustin *et al*., 2009). Liver, adipose tissue and gastrocnemius muscle samples collected from 6 month-old offspring (*n* = 8 control and 9 EtOH; 1 female and/or 1 male per litter) were homogenised and total RNA was extracted using RNeasy Mini Kits (liver; Qiagen, Chadstone, VIC, Australia) or Trizol reagent (muscle and adipose tissue; ThermoFisher Scientific, Richlands, QLD, Australia) according to the manufacturer’s instructions. Total RNA (1 μg/reaction for liver and adipose tissue; 500 ng/reaction for muscle) was reverse transcribed into cDNA using iScript Reverse Transcription Supermix for RT-qPCR (Bio-Rad, Gladesville, NSW, Australia). qPCR reactions were performed on the Applied Biosystems Quantstudio 6 Flex Real-Time PCR System (ThermoFisher Scientific) using 10 ng and 20 ng of cDNA for muscle and liver/adipose tissue respectively, Quantinova Probe PCR Master Mix (Qiagen) and Assay-on-Demand primer/probe sets (see Table 1 for details). Reactions were multiplexed with beta-actin (*Actb*) as an endogenous control. In addition, *Eif2a* and *Rpl19* were used as additional endogenous control genes for liver and muscle/adipose tissue samples respectively (see Table 1). All control genes were stably expressed across all samples, irrespective of experimental group or sex (data not shown). The geometric mean of the two endogenous control genes was used in the ΔΔCt calculation of gene expression, with fold-change expressed relative to the average of the male offspring saline control group.

**Table 1:** Primers used for real-time quantitative PCR analysis of metabolic gene expression. All primers were Assay-on-Demand primer/probe sets from ThermoFisher Scientific (Richlands, QLD, Australia).

| Gene name | Gene symbol | Assay ID | Amplicon size (bp) | Accession number(s) |
| --- | --- | --- | --- | --- |
| Beta actin* | *Actb* | Cat #4351319 | 61 | NM_031144.3 |
| Eukaryotic translation initiation factor 2A* | *Eif2a* | Rn01494813_m1 | 82 | NM_001109339.1 |
| Forkhead box O1 | *Foxo1* | Rn01494868_m1 | 99 | NM_001191846.2 |
| Glucose-6-phosphatase, catalytic subunit | *G6pc* | Rn00689876_m1 | 64 | NM_013098.2 |
| Glucokinase | *Gck* | Rn00561265_m1 | 58 | NM_001270849.1<br>NM_001270850.1<br>NM_012565.2 |
| Glucose transporter 2 [solute carrier family 2 (facilitated glucose transporter), member 2] | *Glut2*<br>(*Slc2a2*) | Rn00563565_m1 | 76 | NM_012879.2 |
| Glucose transporter 4 | *Glut4* | Rn00562597_m1 | 75 | NM_012751.1 |
| Glycogen synthase kinase 3 beta | *Gsk3β* | Rn_00583429_m1 | 71 | NM_032080.1 |
| Insulin receptor | *Insr* | Rn01403321_m1 | 76 | NM_017071.2 |
| Phosphoenolpyruvate carboxykinase 1 | *Pck1* | Rn01529014_m1 | 87 | NM_198780.3 |
| Peroxisome proliferator-activated receptor gamma, coactivator 1 alpha | *Ppargc1a* | Rn00580241_m1 | 94 | NM_031347.1 |
| Ribosomal protein L19* | *Rpl19* | Rn00821265_g1 | 57 | NM_031103.1 |

\* Endogenous control.

Total protein was extracted from visceral adipose tissue (100 mg), gastrocnemius muscle (30 mg) and liver (100-200 mg) samples (1 male offspring only from each litter; *n* = 7 control and 7 EtOH) for Western blot analysis of total and phosphorylated AKT, which plays a central role in insulin signalling in hepatic and peripheral tissues (Mackenzie & Elliott, 2014). Tissues were homogenised in Cell Lysis Buffer (Cell Signaling Technology, Danvers, MA, USA) with protease (Sigma-Aldrich, Sydney, NSW, Australia) and phosphatase (Roche) inhibitors using a FastPrep-24 5G homogeniser (MP Biomedicals, Seven Hills, NSW, Australia). Homogenates were centrifuged (11,000 rpm, 10 min, 4°C) and resultant supernatant assayed using a DC Protein Assay Kit (Bio-Rad). A total of 20 μg of sample was loaded per well onto 12% SDS-PAGE gels and subsequently transferred overnight at 4°C to Immun-Blot LF PVDF membranes (Bio-Rad). Membranes were incubated overnight with one of the following rabbit primary antibodies (all from Cell Signaling Technology and all cross-react with rat): anti-Pan-AKT (1:1000, Cat# 5373s, 60 kDa, RRID:AB_10891424), anti-phospho-AKT_Thr308_ (1:1000, Cat# 9275s, 60 kDa, RRID:AB_329828), anti-phospho-AKT_Ser473_ (1:1000, Cat# 4060s, 60 kDa, RRID:AB_2315049). Protein expression was measured as previously described using a LI-COR Odyssey CLx infrared imaging system (LI-COR Biosciences, Lincoln, NE, USA, RRID:SCR_014579) following exposure to LI-COR IRDye 680 goat anti-rabbit secondary antibody (LI-COR Biosciences, Cat# 926-32221, RRID:AB_621841). Band intensity was analysed using Image Studio Lite software (LI-COR Biosciences, v5.2, RRID:SCR_013715) and was normalised to glyceraldehyde 3-phosphate dehydrogenase (GAPDH, 1:1000, Cat# 2118s, 37 kDa, RRID:AB_561053) or β-actin (ACTB, 1:1000, Cat#4970, 45 kDa, RRID:AB_2223172) immunoreactivity for densitometric analysis. Bands were quantified by an assessor blinded to treatment. Protein levels in the EtOH samples were expressed relative to the average of the saline controls.

### Plasma insulin, glucose and lipid analysis

Fasting plasma samples collected from offspring at PN30 and during the GTT at 6-7 months of age were analysed for insulin levels using a rat insulin radioimmunoassay kit (SRI-13K, Millipore Australia, Kilsyth, VIC). Insulin samples were run in duplicate at 1:5, 1:10 or 1:20 dilution. Assay sensitivity was 0.03 ng/mL and inter- and intra-assay coefficients of variation were 16.7% and 12.1% respectively. Plasma collected at offspring culls at PN30 were analysed for triglycerides (TG), high-density lipoproteins (HDL), and low-density lipoproteins (LDL) using a Cobas Integra 400 Plus Chemistry Analyzer (Block Scientific, Bellport, NY, USA). In addition, plasma collected at baseline for the GTT at 6 months of age were analysed for glucose levels using the Cobas Analyzer, for comparison to measurement of glucose levels by the glucometer. Blood glucose levels, as measured using the glucometer, were found to underestimate plasma glucose levels measured using the ‘gold standard’ Cobas analyser (Figure 2A). However, this underestimation was consistent across all treatment/sex combinations (Figure 2B). Therefore, where insufficient plasma was available for Cobas analysis (PN30), glucometer readings of blood glucose levels were adequate to compare fasting glucose and insulin sensitivity/resistance indices between treatment/sex groups. When sufficient plasma could be collected (6 months of age), plasma glucose levels were used for all analyses.

**Figure 2:**
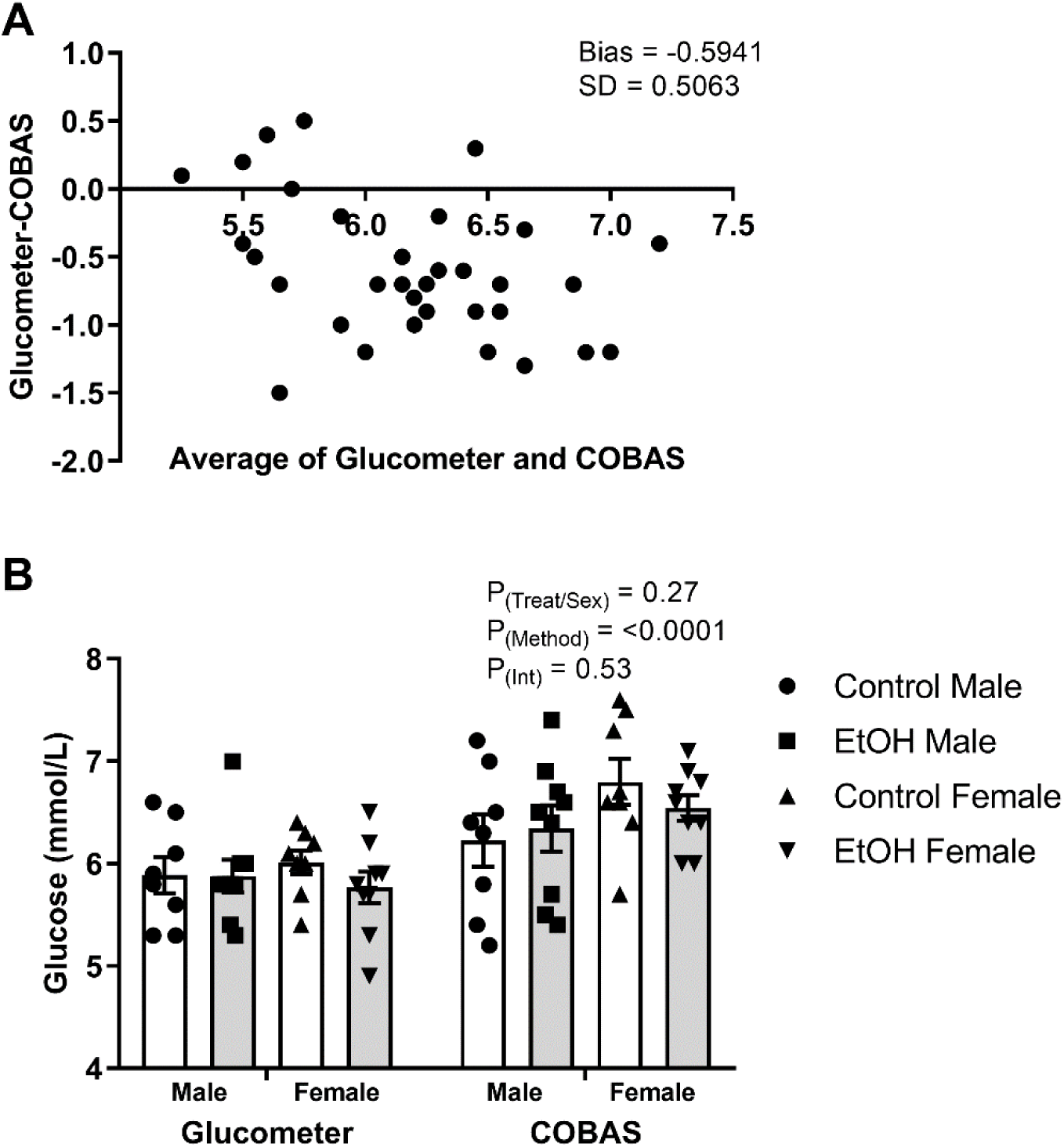
Comparison of blood glucose concentrations measured by glucometer and plasma glucose concentrations measured by Cobas analyser. Dual-analysis was conducted on *n* = 8 control litters and *n* = 9 ethanol litters, with 1 male and 1 female analysed per litter. (A) Bland-Altman plot to visualise the difference in the glucometer and Cobas readings versus the average of the readings. (B) Comparison of glucose levels across each treatment/sex group by method. White bars indicate control and black bars indicate ethanol-exposed (EtOH) animals within each sex. Data analysed using a two-way ANOVA, with treatment/sex (Control Male, EtOH Male, Control Female, EtOH Female) and method (glucometer/Cobas) as factors.

### Calculations and statistical analyses

Area under the glucose (AUGC) and insulin (AUIC) curves were calculated using the trapezius method with baseline defined as the basal level prior to injection (Allison *et al*., 1995). Glucose concentration curves for the ITT were inverted before calculating AUGC as previously described (Gardebjer *et al*., 2015; Nagy & Einwallner, 2018). Acute 1^st^ phase insulin secretion was calculated as the AUIC from basal to 5 min post-glucose bolus and 2^nd^ phase insulin secretion as the AUIC from 5 to 90 min post-bolus. Note that baseline was set as zero for the 2^nd^ phase secretion analysis, given that 5 min post-bolus was not at basal levels. Quantitative insulin sensitivity check index (QUICKI) was calculated using 1/[log(fasting insulin (μU/mL)) + log(fasting glucose (mg/dL)] (Katz *et al*., 2000). The homeostasis model assessment-estimated insulin resistance (HOMA-IR) was calculated using [fasting insulin (μU/mL) × fasting glucose (mg/dL)]/2430, which has been validated for use in rats (Cacho *et al*., 2008). Total cholesterol (TC) was calculated by re-arranging the Friedewald equation (Friedewald *et al*., 1972) such that TC = HDL + LDL + (TG/5).

Analyses were conducted using GraphPad Prism 7.0 (GraphPad Software, San Diego, CA, USA) and all data presented as mean ± SEM. Prior to analysis, all data were tested for normality using the D’Agnostino-Pearson omnibus or the Shapiro-Wilk normality tests. Maternal parameters and Western blot data were analysed using a Student’s t-test (parametric data) or a Mann-Whitney U-test (non-parametric data). Where variances were significantly different but data was normally distributed, Welch’s correction was applied. All other data were analysed using a two-way ANOVA, with offspring sex (male/female) and treatment group (EtOH/Control) as factors. Where there was a significant interaction, a Sidak multiple comparison test was used to determine significantly different groups. Where data was not normally distributed, a non-parametric Kruskal-Wallis test across all treatment/sex groups was used, with Dunn’s multiple comparison test used to determine significantly different groups but only treatment differences within sex or sex differences with treatment are reported. Significance level was *P*<0.05 for all statistical tests, with 0.05 ≥ *P* ≤ 1.0 considered indicative of trends.

## Results

### Maternal parameters and postnatal growth

Dams in the EtOH and Control groups were of similar weight at mating and at E13.5 (Table 2). Gestational weight gain prior to gavage and post-gavage were also not significantly different between groups, nor were daily chow or water consumption (Table 2). When energy intake contributed from chow consumption and the EtOH treatment (∼9-10 kJ) on the days of gavage were calculated, there was no difference in energy intake between the EtOH and Control dams (Table 2). Blood glucose concentrations at 1 h and 5 h post gavage on E13.5 and E14.5 were not significantly different (Table 2). There were also no differences in pregnancy outcomes between EtOH and Control dams, such as litter sex ratio, litter size, and number of implantation scars (Table 2). However, one dam in the EtOH group had a litter of only 7 pups due to a suspected blockage in one uterine horn (no implantation scars were observed) and so this litter was excluded from subsequent analyses. BAC was measured in 4 out of 8 Control dams, with BAC below the limit of detection at each time point. In EtOH-treated dams, mean BAC was ∼50 mg/dL (∼0.05%) at 1 h following gavage at E13.5 and E14.5, but by 5 h post-gavage, was below the limit of detection on both days (Table 2).

**Table 2:** Maternal parameters for dams treated with saline (Control) or ethanol (EtOH). All animals treated by oral gavage on embryonic day 13.5 and 14.5.

| Maternal parameters | Treatment group |  | *P* value |
| --- | --- | --- | --- |
|  | Control<br>(*n* = 8) | EtOH<br>(*n* = 10) |  |
| Body weight when mated (g) | 251 ± 4 | 255 ± 4 | 0.61 |
| Body weight at E13.5 (g) | 310 ± 5 | 312 ± 5 | 0.77 |
| Total weight gain (g)* | 152 ± 9 | 148 ± 5 | 0.98 |
| Pre-gavage (mating to E12.5) | 59 ± 3 | 58 ± 4 | 0.85 |
| Post-gavage (E15.5 to birth) | 91 ± 6 | 89 ± 5 | 0.77 |
| H<sub>2</sub>O consumption (mL/day) |  |  |  |
| Pre-gavage (E12.5) | 31.9 ± 1.7 | 29.6 ± 1.8 | 0.37 |
| During gavage (E13.5-E14.5) | 25.8 ± 1.3 | 25.3 ± 1.4 | 0.79 |
| Post-gavage (E15.5 to birth) | 36.9 ± 1.1 | 34.8 ± 1.6 | 0.33 |
| Chow consumption (g/day) |  |  |  |
| Pre-gavage (E12.5) | 24.7 ± 1.2 | 24.4 ± 0.8 | 0.79 |
| During gavage (E13.5-E14.5) | 20.9 ± 1.0 | 20.4 ± 1.0 | 0.70 |
| Post-gavage (E15.5 to birth) | 26.6 ± 0.7 | 26.0 ± 0.9 | 0.61 |
| Energy consumption (kJ/day) |  |  |  |
| E13.5* | 299.3 ± 14.6 | 283.4 ± 6.2 | 0.70 |
| E14.5 | 286.0 ± 18.8 | 305.0 ± 24.0 | 0.56 |
| Blood glucose (mmol/L) |  |  |  |
| 1h post E13.5 gavage* | 5.8 ± 0.2 | 6.1 ± 0.2 | 0.13 |
| 5h post E13.5 gavage | 5.9 ± 0.1 | 6.1 ± 0.2 | 0.38 |
| 1h post E14.5 gavage | 5.7 ± 0.2 | 6.0 ± 0.1 | 0.09 |
| 5h post E14.5 gavage* | 6.2 ± 0.2 | 6.0 ± 0.1 | 0.62 |
| Litter sex ratio (M:F) | 1.1 ± 0.2 | 1.2 ± 0.2 | 0.78 |
| Litter size | 15<sup>†</sup> | 14<sup>†</sup> | 0.47 |
| Number of implantation scars* | 16<sup>†</sup> | 15 ± 1<sup>†</sup> | 0.99 |
| Blood alcohol concentration (BAC) (mg/dL) |  |  |  |
| 1h post E13.5 gavage | <LD<sup>‡</sup> | 48.2 ± 7.5 | - |
| 5h post E13.5 gavage | <LD<sup>‡</sup> | <LD | - |
| 1h post E14.5 gavage | <LD<sup>‡</sup> | 52.9 ± 4.5 | - |
| 5h post E14.5 gavage | <LD<sup>‡</sup> | <LD | - |

Data are presented as mean ± SEM. *P* values were obtained using an unpaired Student's *t*-test.
<LD = below the limit of detection; E = embryonic day; EtOH = ethanol; M = male; F = female.
\* Indicates data not normally distributed and analysed with a non-parametric Mann-Whitney U-test.
† Denotes parameters expressed as whole pups/scars.
‡ Measured in 4 animals only.

There was no difference in pup weights at PN1 between EtOH and control litters, nor differences in weight gain until weaning at PN21 (Table 3). For the first week post-weaning, weight gain was higher in males than in females (*P_(sex)_* = 0.03) but there was no effect of treatment (Table 3). Weights of offspring at the PN30 cull were also not different (Table 3).

**Table 3:**
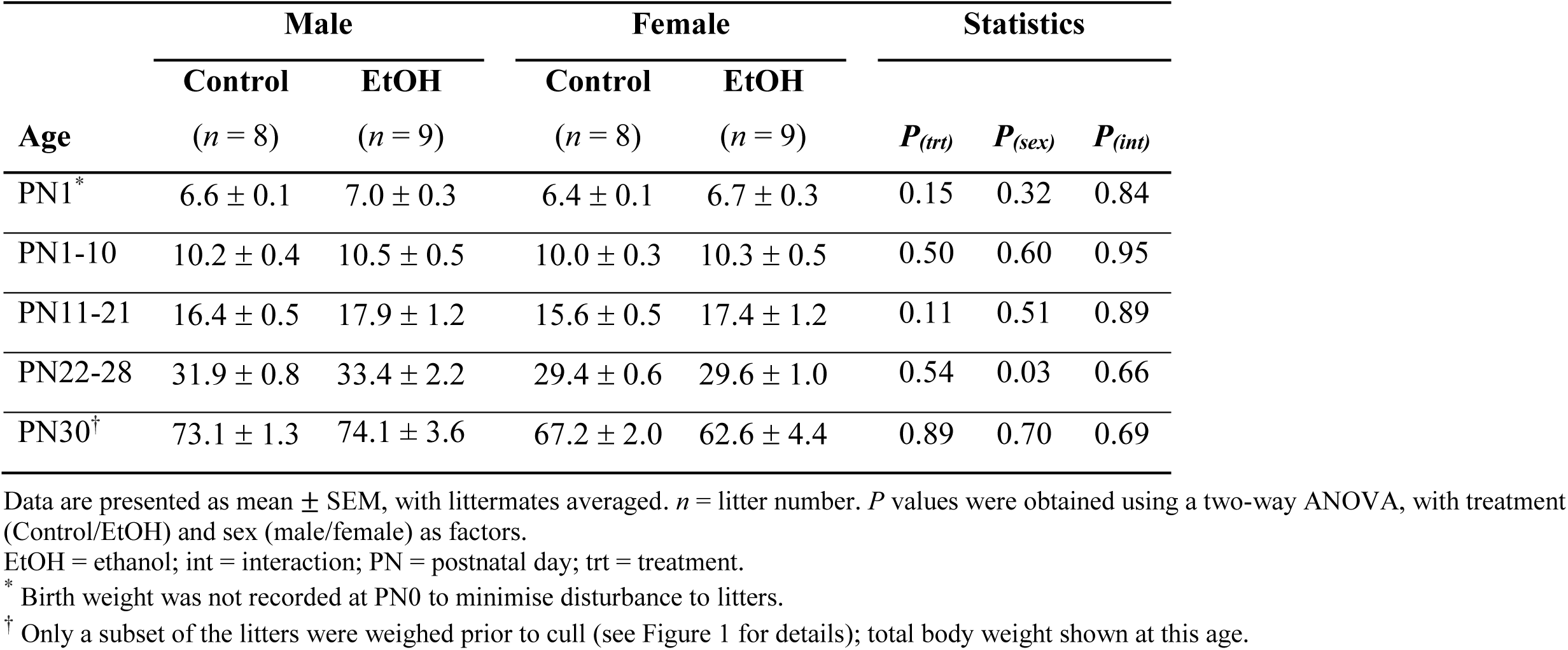
Postnatal weight and growth of offspring from saline (Control) and ethanol (EtOH) treated litters during the first month. All offspring from dams treated by oral gavage at embryonic day 13.5 and 14.5. Weights at PN1 and PN30 are average body weight (g). At other ages, average total weight gain over the specified age ranges are shown (g).

| Age | Male |  | Female |  | Statistics |  |  |
| --- | --- | --- | --- | --- | --- | --- | --- |
|  | Control<br>(*n* = 8) | EtOH<br>(*n* = 9) | Control<br>(*n* = 8) | EtOH<br>(*n* = 9) | *P*<sub>(trt)</sub> | *P*<sub>(sex)</sub> | *P*<sub>(int)</sub> |
| PN1<sup>*</sup> | 6.6 ± 0.1 | 7.0 ± 0.3 | 6.4 ± 0.1 | 6.7 ± 0.3 | 0.15 | 0.32 | 0.84 |
| PN1-10 | 10.2 ± 0.4 | 10.5 ± 0.5 | 10.0 ± 0.3 | 10.3 ± 0.5 | 0.50 | 0.60 | 0.95 |
| PN11-21 | 16.4 ± 0.5 | 17.9 ± 1.2 | 15.6 ± 0.5 | 17.4 ± 1.2 | 0.11 | 0.51 | 0.89 |
| PN22-28 | 31.9 ± 0.8 | 33.4 ± 2.2 | 29.4 ± 0.6 | 29.6 ± 1.0 | 0.54 | 0.03 | 0.66 |
| PN30<sup>†</sup> | 73.1 ± 1.3 | 74.1 ± 3.6 | 67.2 ± 2.0 | 62.6 ± 4.4 | 0.89 | 0.70 | 0.69 |

Data are presented as mean ± SEM, with littermates averaged. *n* = litter number. *P* values were obtained using a two-way ANOVA, with treatment (Control/EtOH) and sex (male/female) as factors.
EtOH = ethanol; int = interaction; PN = postnatal day; trt = treatment.
<sup>\*</sup> Birth weight was not recorded at PN0 to minimise disturbance to litters.
<sup>†</sup> Only a subset of the litters were weighed prior to cull (see Figure 1 for details); total body weight shown at this age.

### Fasting blood glucose, plasma insulin and lipid levels in adolescent offspring

Fasting blood glucose, plasma insulin and indexes for insulin resistance (HOMA-IR) and insulin sensitivity (QUICKI) were not significantly different between groups in either sex at PN30 (Table 4). Fasting blood glucose (mmol/L) was also not different at 10-11 weeks of age (Male EtOH: 5.5 ± 0.2, *n* = 9; Male Control: 5.5 ± 0.2, *n* = 8; Female EtOH: 5.3 ± 0.1, n = 9; Female Control: 5.5 ± 0.1, *n* = 8. *P*_treat_ = 0.69, *P*_sex_ = 0.51, *P*_int_ = 0.57). The plasma lipid profile (HDL, LDL, TG and TC) was also not different between EtOH and control offspring at PN30 (Table 4).

**Table 4:** Effect of prenatal ethanol exposure on fasting blood glucose, plasma insulin and plasma lipid levels in adolescent offspring. Offspring were prenatally exposed to ethanol (EtOH) or saline (control) at embryonic day 13.5 and 14.5. All parameters were measured at postnatal day 30.

| Parameters | Male |  | Females |  | Statistics |  |  |
| --- | --- | --- | --- | --- | --- | --- | --- |
|  | Control<br>(*n* = 6) | EtOH<br>(*n* = 8) | Control<br>(*n* = 3) | EtOH<br>(*n* = 3) | *P*<sub>(trt)</sub> | *P*<sub>(sex)</sub> | *P*<sub>(int)</sub> |
| Fasting blood glucose (mmol/L) | 5.6 ± 0.2 | 5.9 ± 0.2 | 6.1 ± 0.2 | 5.9 ± 0.1 | 0.64 | 0.69 | 0.83 |
| Fasting plasma insulin (ng/mL) | 0.55 ± 0.07 | 0.65 ± 0.06 | 0.64 ± 0.11 | 0.63 ± 0.07 | 0.58 | 0.67 | 0.53 |
| HOMA-IR | 0.71 ± 0.11 | 0.81 ± 0.07 | 0.83 ± 0.15 | 0.79 ± 0.07 | 0.78 | 0.65 | 0.54 |
| QUICKI | 0.31 ± 0.007 | 0.30 ± 0.003 | 0.30 ± 0.007 | 0.30 ± 0.003 | 0.55 | 0.50 | 0.50 |
| Plasma HDL (mmol/L) | 1.07 ± 0.15 | 0.98 ± 0.21 | 0.87 ± 0.16 | 0.80 ± 0.18 | 0.73 | 0.41 | 0.95 |
| Plasma LDL (mmol/L) | 0.43 ± 0.06 | 0.46 ± 0.04 | 0.52 ± 0.07 | 0.33 ± 0.01 | 0.26 | 0.80 | 0.11 |
| Plasma triglycerides (mmol/L) | 0.92 ± 0.10 | 1.09 ± 0.21 | 0.83 ± 0.20 | 0.78 ± 0.14 | 0.53 | 0.22 | 0.99 |
| Plasma total cholesterol (mmol/L)* | 1.68 ± 0.18 | 1.65 ± 0.23 | 1.55 ± 0.20 | 1.29 ± 0.16 | 0.59 | 0.35 | 0.65 |

EtOH = ethanol; HDL = high density lipoprotein; HOMA-IR = homeostatic model assessment of insulin resistance; int = interaction; LDL = low density lipoprotein; QUICKI = quantitative insulin-sensitivity check index; trt = treatment.
\* Calculated using HDL + LDL + (triglycerides/5).

### Food preference and water consumption at 4-5 months of age

There were no significant differences in consumption of SD or HFD between EtOH and Control offspring at any time (Table 5). However, as the HFD was more energy dense than the SD (19.4 MJ/kg versus 14.0 MJ/kg), there was a significant decrease in energy intake per day by the EtOH groups compared to controls, due to a slightly reduced (non-significant) consumption of HFD by these groups. Animals consumed ∼10-fold more HFD than SD during CP1, with a concomitant 10-fold decrease in SD consumption compared to baseline levels. This continued through CP2, although HFD consumption dropped to ∼4-fold that of SD. However, there were sex differences in consumption rates, with females consuming more SD at baseline and more HFD during CP1 and CP2 than males (Table 5).

**Table 5:** Offspring food and water consumption throughout the food preference study. All animals were housed individually and acclimatised to the study cage for 4 days. Following acclimatisation, baseline measurements of standard chow diet (SD) and water were conducted over days 1-4. A choice of high fat diet (HFD) or SD was then offered for the next 4 days, divided into choice period (CP)1 (days 5-6) and CP2 (days 7-8).

| Parameter | Male |  | Female |  | Statistics |  |  |
| --- | --- | --- | --- | --- | --- | --- | --- |
| | Control | EtOH | Control | EtOH | $P_{(trt)}$ | $P_{(sex)}$ | $P_{(int)}$ |
| | ($n = 8$) | ($n = 9$) | ($n = 8$) | ($n = 9$) | | | |
| ***H<sub>2</sub>O consumption (mL/g BW/day)*** |  |  |  |  |  |  |  |
| Baseline | 0.064 ± 0.006 | 0.060 ± 0.002 | 0.070 ± 0.005 | 0.082 ± 0.005 | 0.34 | 0.007 | 0.12 |
| CP1 | 0.049 ± 0.004 | 0.047 ± 0.002 | 0.058 ± 0.011 | 0.068 ± 0.004 | 0.37 | <0.001 | 0.19 |
| CP2* | 0.046 ± 0.003 | 0.049 ± 0.003 | 0.055 ± 0.003 | 0.070 ± 0.004<sup>†</sup> | $P = 0.0004$ | | |
| ***Standard chow diet (SD) consumption (g/g BW/day)*** |  |  |  |  |  |  |  |
| Baseline | 0.047 ± 0.001 | 0.046 ± 0.001 | 0.053 ± 0.002 | 0.054 ± 0.001 | 0.99 | <0.001 | 0.51 |
| CP1 | 0.007 ± 0.003 | 0.005 ± 0.001 | 0.008 ± 0.004 | 0.003 ± 0.003 | 0.25 | 0.92 | 0.63 |
| CP2 | 0.019 ± 0.005 | 0.017 ± 0.003 | 0.010 ± 0.004 | 0.013 ± 0.003 | 0.87 | 0.11 | 0.44 |
| ***High fat diet (HFD) consumption (g/g BW/day)*** |  |  |  |  |  |  |  |
| CP1 | 0.055 ± 0.004 | 0.055 ± 0.003 | 0.077 ± 0.006 | 0.072 ± 0.004 | 0.54 | <0.0001 | 0.64 |
| CP2 | 0.046 ± 0.004 | 0.043 ± 0.004 | 0.067 ± 0.004 | 0.060 ± 0.004 | 0.27 | <0.0001 | 0.61 |
| ***Energy intake (kJ/day)*** |  |  |  |  |  |  |  |
| Baseline | 357.8 ± 8.4 | 348.3 ± 10.3 | 230.4 ± 10.4 | 222.1 ± 6.5 | 0.33 | <0.0001 | 0.95 |
| CP1 | 628.3 ± 20.6 | 605.8 ± 26.9 | 502.2 ± 27.1 | 430.4 ± 20.1 | 0.06 | <0.0001 | 0.31 |
| CP2 | 579.8 ± 25.6 | 538.5 ± 26.6 | 457.1 ± 22.5 | 396.0 ± 17.6 | 0.04 | <0.0001 | 0.67 |

Data are presented as mean ± SEM, with 1 male and 1 female used per litter ( $n$ = individual offspring). $P$ values were obtained using a two-way ANOVA, with treatment (Control/EtOH) and sex (male/female) as factors.
CP = choice period; HFD = high-fat diet; BW = body weight.
\* Indicates data not normally distributed and analysed with a non-parametric Kruskal-Wallis Test across all 4 groups.
† Indicates significantly different to corresponding treatment group in males ( $P=0.006$ ) from Dunn's multiple comparison procedure.

Daily baseline water and SD consumption was not significantly different between Control and EtOH offspring in either sex, but females did drink a larger volume of water compared to males, regardless of treatment (Table 5). EtOH females also increased their water consumption compared to controls in the 2^nd^ half of the testing period (CP2, Table 5), with a non-parametric Mann-Whitney U-test performed separately for each sex indicating a significant difference between control and EtOH females (*P* = 0.003) but no difference between groups within males (*P* = 0.34).

### Fasting plasma glucose, glucose clearance and insulin sensitivity in adult offspring

At 6 months of age, fasting plasma glucose continued to be similar between EtOH and Control groups and between males and females (Figure 3A). However, fasting plasma insulin was significantly elevated in ETOH-exposed males compared to control males, with females showing no effect of treatment (Figure 3B). HOMA-IR and QUICKI indices were also significantly altered in EtOH compared to Control offspring, indicative of increased insulin resistance and reduced insulin sensitivity (Figure 3C, D).

**Figure 3:**
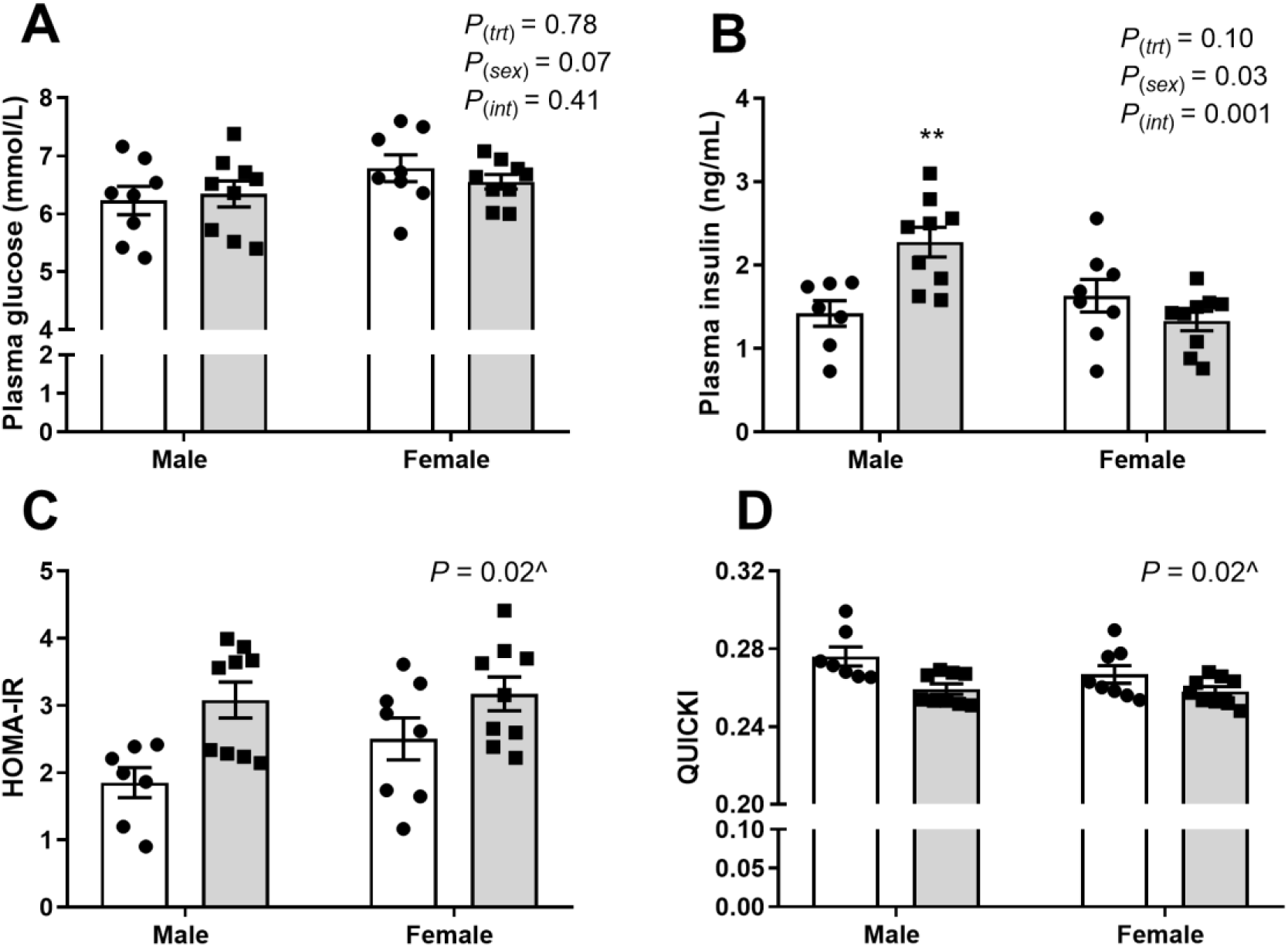
Effect of prenatal alcohol exposure on fasting plasma glucose and plasma insulin levels in adult offspring. Offspring were prenatally exposed to ethanol (grey) or saline control (white). (A) Fasting plasma glucose and (B) plasma insulin were measured at 6 months of age. Corresponding homeostatic model assessment of insulin resistance (HOMA-IR) (C) and quantitative insulin-sensitivity check index (QUICKI) (D) were calculated. Data are presented as mean ± SEM. *n* = 8 control litters and *n* = 9 ethanol litters, with 1 male and 1 female analysed per litter. Data for A) and B) were analysed with two-way ANOVA; while data for C) and D) were analysed using a non-parametric Kruskal-Wallis test due to non-normal data distribution (indicated by ^). ** = significantly different to control male by Sidak’s multiple comparison post-hoc procedure.

In the GTT, there was no significant difference in the response to glucose injection (as shown in Figure 4A) and overall AUGC between EtOH and Control offspring (Figure 4B). Insulin output, as shown by the AUIC, was significantly elevated in males compared to females during the GTT (Figure 4C, D). This was particularly evident during the acute first-phase insulin response, with EtOH males having a tendency to increase insulin output compared to Control male and female offspring, although this did not reach statistical significance following the non-parametric Dunn’s multiple comparison procedure (Figure 4E). Second phase insulin production continued to be higher in males than females, but there was no effect of PAE (Figure 4F). In addition, there were significant differences in glucose clearance in response to exogenous insulin in the ITT (Figure 4G, H). EtOH male offspring had significantly reduced AUGC compared to EtOH females by Dunn’s multiple comparison procedure, but this group was also significantly different to control male offspring using a non-parametric Mann-Whitney test (*P* = 0.0079). However, PAE had no effect on glucose clearance in females during the ITT (Figure 4H).

**Figure 4:**
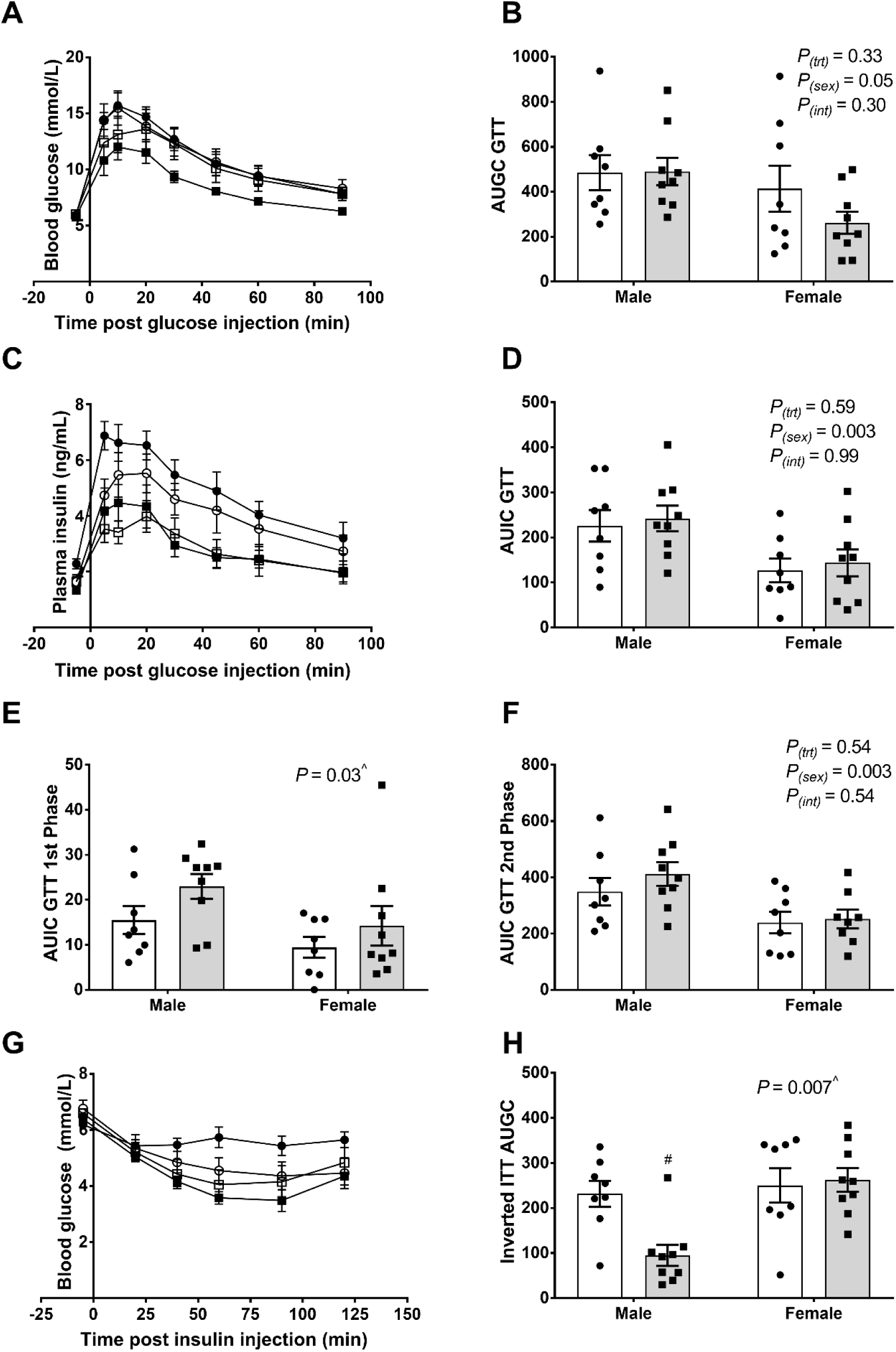
Effect of prenatal alcohol exposure on glucose clearing and insulin sensitivity. Offspring were prenatally exposed to ethanol (grey) or saline (white). (A) Blood glucose concentration curve during a glucose tolerance test (GTT); (B) area under the glucose curve (AUGC) generated from the GTT; (C) plasma insulin secretion during the GTT; (D) area under the insulin curve (AUIC) generated from the GTT; (E) AUIC for 1^st^ phase insulin secretion; and (F) AUIC for 2^nd^ phase insulin secretion. (G) Blood glucose concentrations following an insulin tolerance test (ITT) and (H) AUGC from inverted ITT curves. Control males (white circles), ethanol males (black circles), control females (white squares) and ethanol females (black squares) (A, C, G). Data represented as mean ± SEM; *n* = 8 control litters and *n* = 9 ethanol litters, with 1 male and 1 female analysed per litter. Data were analysed with two-way ANOVA (B, D, F) or a non-parametric Kruskal-Wallis test (indicated by ^) due to non-normal data distribution (E, H). # indicates significant difference between sexes within treatment (*P*<0.05), where significance was determined by Dunn’s multiple comparisons test.

### Body condition and organ weights at 6 months of age

At the completion of the study, all animals were culled and a subset (1-2 males and/or females per litter for *n*=12 per sex per group) underwent detailed morphometric analysis. There was no significant difference in body weight, measures of size (snout-rump length, tibia length), or measures of body condition (ponderal index, absolute abdominal circumference and abdominal circumference relative to body size parameters) between EtOH-exposed and control offspring (Table 6). However, as would be expected, males were significantly larger than females across all parameters (Table 6). There was also no significant difference in liver or pancreas weight (absolute or relative to body weight) between EtOH and control offspring. Again, liver weight (absolute and relative to BW) and absolute pancreas weight were significantly higher in males than females. However, pancreas weight relative to body weight was significantly higher in females than males (Table 6).

**Table 6:**
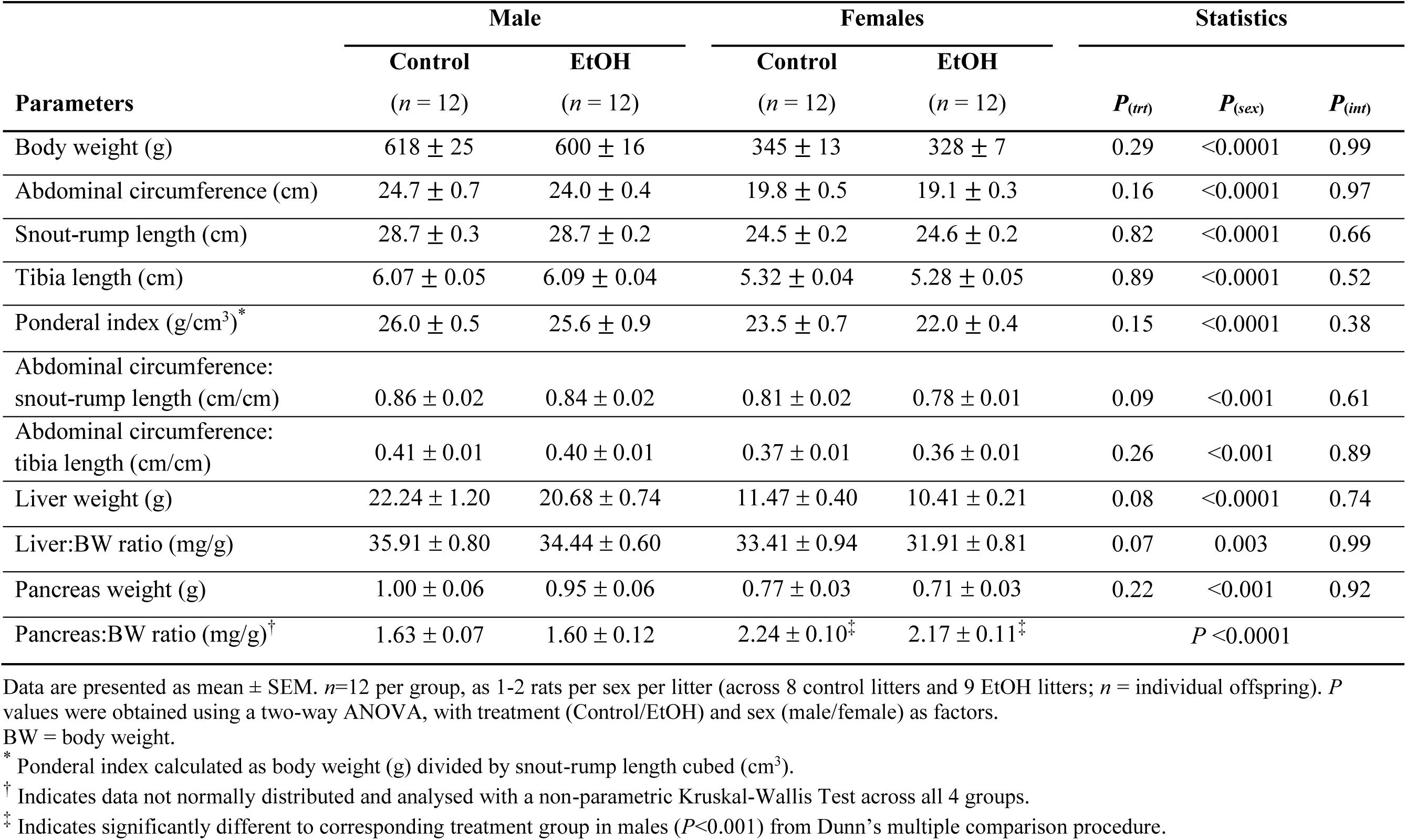
Body condition and organ weights of offspring from saline (Control) and ethanol (EtOH) treated litters at 6 months of age. Offspring were prenatally exposed to ethanol (EtOH) or saline (control) at embryonic day 13.5 and 14.5.

### Expression of hepatic genes involved in insulin signalling and glucose homeostasis

Liver tissue was collected from 6 month-old offspring for analysis of genes involved in glucose transport (*Glut2*), homeostatic regulation of glucose levels (*G6pc*, *Gck*, *Pck1* and *Ppargc1a*) and insulin signalling (*Insr*) (Table 7). There was no significant difference in the expression of these genes between EtOH-exposed and control offspring (Table 7). However, there was a sex-specific difference in expression of *Ppargc1a* and *Insr*, and a trend for an effect of sex on *Glut2* expression, with all three genes displaying higher relative expression in females compared with males (Table 7).

**Table 7:**
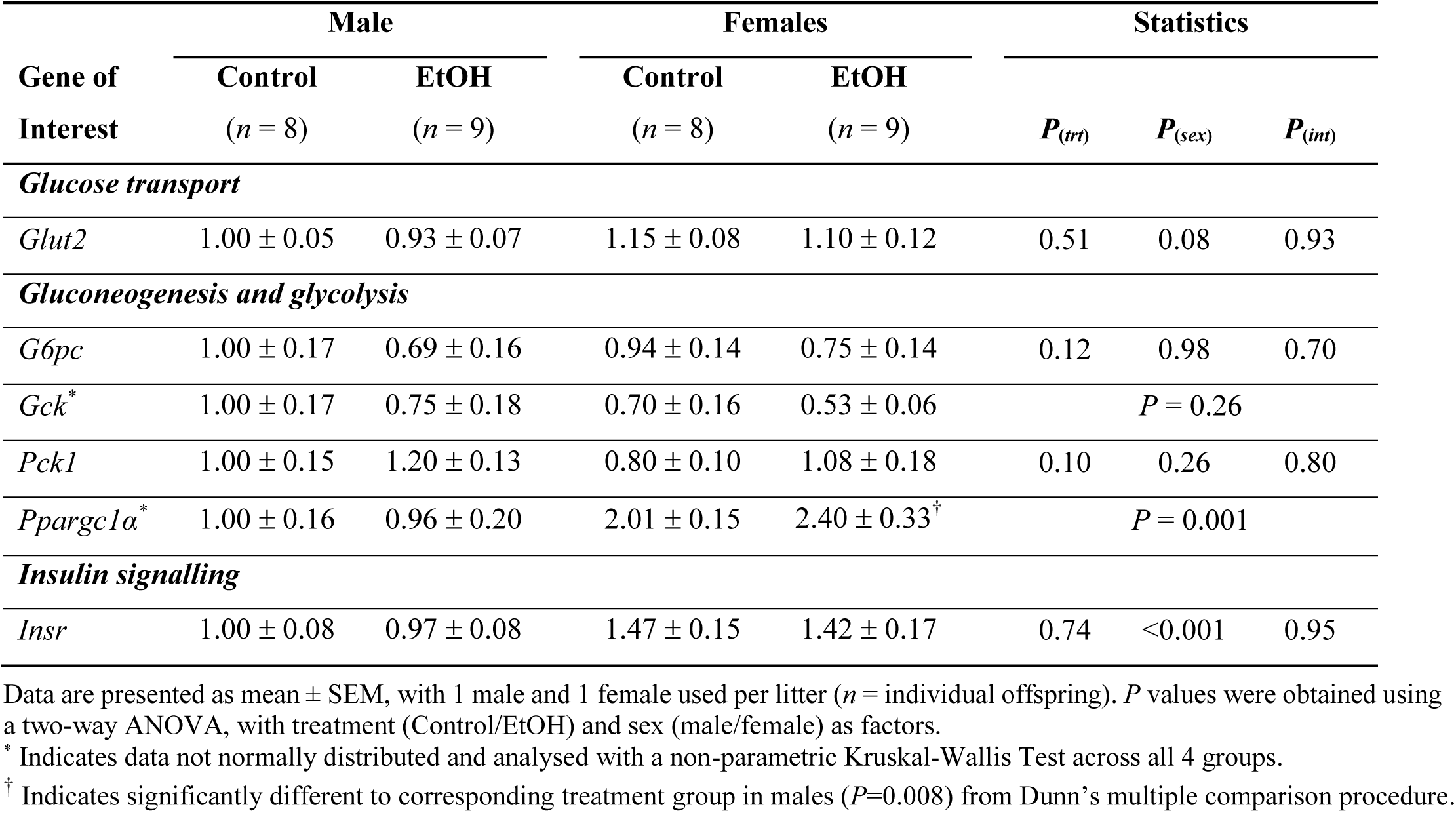
Expression of hepatic genes involved in glucose and insulin signalling. Offspring were prenatally exposed to ethanol (EtOH) or saline (control) at embryonic day 13.5 and 14.5. Liver tissue was collected at 6 months of age.

| Gene of Interest | Male |  | Females |  | Statistics |  |  |
| --- | --- | --- | --- | --- | --- | --- | --- |
| | Control | EtOH | Control | EtOH | $P_{(trt)}$ | $P_{(sex)}$ | $P_{(int)}$ |
| | ($n = 8$) | ($n = 9$) | ($n = 8$) | ($n = 9$) | | | |
| *Glucose transport* |  |  |  |  |  |  |  |
| *Glut2* | 1.00 ± 0.05 | 0.93 ± 0.07 | 1.15 ± 0.08 | 1.10 ± 0.12 | 0.51 | 0.08 | 0.93 |
| *Gluconeogenesis and glycolysis* |  |  |  |  |  |  |  |
| *G6pc* | 1.00 ± 0.17 | 0.69 ± 0.16 | 0.94 ± 0.14 | 0.75 ± 0.14 | 0.12 | 0.98 | 0.70 |
| *Gck* * | 1.00 ± 0.17 | 0.75 ± 0.18 | 0.70 ± 0.16 | 0.53 ± 0.06 | $P = 0.26$ | | |
| *Pck1* | 1.00 ± 0.15 | 1.20 ± 0.13 | 0.80 ± 0.10 | 1.08 ± 0.18 | 0.10 | 0.26 | 0.80 |
| *Ppargc1α* * | 1.00 ± 0.16 | 0.96 ± 0.20 | 2.01 ± 0.15 | 2.40 ± 0.33<sup>†</sup> | $P = 0.001$ | | |
| *Insulin signalling* |  |  |  |  |  |  |  |
| *Insr* | 1.00 ± 0.08 | 0.97 ± 0.08 | 1.47 ± 0.15 | 1.42 ± 0.17 | 0.74 | <0.001 | 0.95 |

Data are presented as mean ± SEM, with 1 male and 1 female used per litter (*n* = individual offspring). *P* values were obtained using a two-way ANOVA, with treatment (Control/EtOH) and sex (male/female) as factors.
\* Indicates data not normally distributed and analysed with a non-parametric Kruskal-Wallis Test across all 4 groups.
<sup>†</sup> Indicates significantly different to corresponding treatment group in males (*P*=0.008) from Dunn's multiple comparison procedure.

### Hepatic and peripheral tissue insulin signalling

Given that insulin resistance was only observed in EtOH-exposed males compared to control males, and there was no effect of EtOH exposure on females, we focused our molecular analysis of insulin signalling on male offspring. We examined a key molecule involved in the insulin signalling pathway both centrally and in peripheral tissues, AKT, also known as protein kinase B. In liver (Figure 5A-E) and skeletal muscle (Figure 5F-J), there was no significant difference in total (pan-)AKT, pAKT_Ser473_ (absolute or as a ratio of pan-AKT) or pAKT_Thr308_ (absolute or as a ratio of pan-AKT) between EtOH-exposed and control male offspring. Muscle gene expression of the glucose transporter, *Glut4*, and the insulin receptor, *Insr*, were also not affected by EtOH exposure (Table 8). However, there was a significant increase in pan-AKT in adipose tissue from EtOH-exposed males compared to controls (Figure 5K) and a tendency for increased pAKT_Thr308_ (Figure 5M, *P* = 0.07), but not as a ratio of pan-AKT (Figure 5O). Expression of genes involved in glucose transport (*Glut 4*) and insulin signalling both upstream (*Insr*) and downstream (*Foxo1*, *Gsk3β*) of AKT were not altered in adipose tissue from EtOH-exposed male offspring (Table 8).

**Figure 5:**
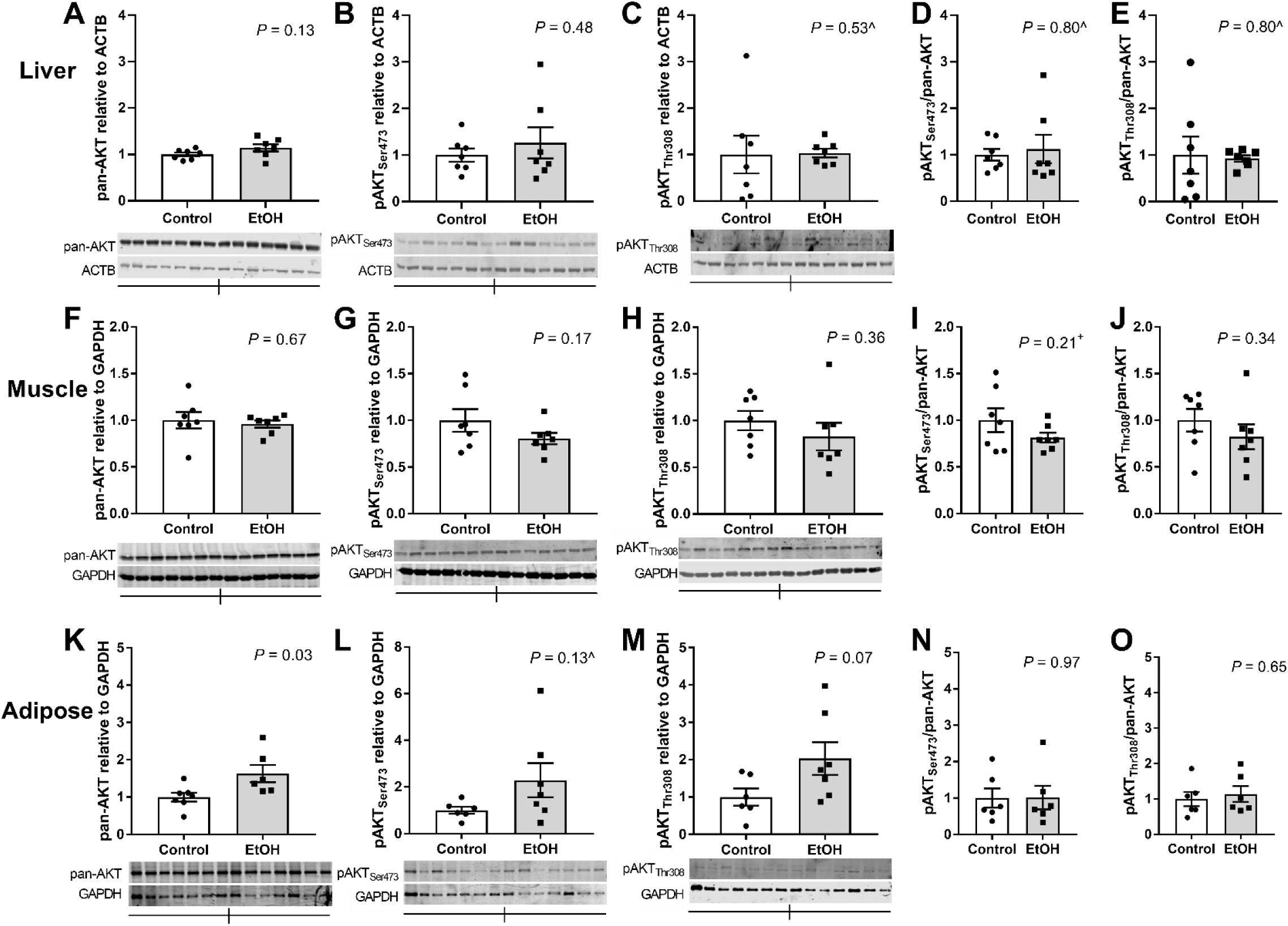
Effect of prenatal alcohol on AKT protein levels in liver and peripheral tissues in male offspring at 6 months of age. Protein levels and phosphorylation state, expressed relative to ACTB, in liver (A-E) and expressed relative to GAPDH in gastrocnemius muscle (F-J) and adipose tissue (K-O). Offspring were prenatally exposed to ethanol (grey) or saline (white). Data represented as mean ± SEM. *n* = 7 litters per group with 1 male per litter. Data were analysed with an unpaired t-test (normally distributed data) or a Mann-Whitney U-test (non-normal distribution, indicated by ^). ^+^ indicates Welch’s correction was applied due to unequal variances. ACTB = β-actin; GAPDH = glyceraldehyde 3-phosphate dehydrogenase.

**Table 8:** Expression of genes involved in glucose transport and insulin signalling in peripheral tissues. Offspring were prenatally exposed to ethanol (EtOH) or saline (control) at embryonic day 13.5 and 14.5. Gastrocnemius muscle and adipose tissue were collected at 6 months of age.

| Gene of Interest | Treatment group |  | *P* value |
| --- | --- | --- | --- |
|  | Control<br>(*n* = 8) | EtOH<br>(*n* = 9) |  |
| *Gastrocnemius muscle* |  |  |  |
| *Glut4* | 1.00 ± 0.30 | 1.11 ± 0.33 | 0.81 |
| *Insr*<sup>*</sup> | 1.00 ± 0.16 | 0.99 ± 0.72 | 0.38 |
| *Adipose tissue* |  |  |  |
| *Glut4* | 1.00 ± 0.11 | 0.85 ± 0.12<sup>†</sup> | 0.38 |
| *Insr* | 1.00 ± 0.08 | 0.83 ± 0.08<sup>†</sup> | 0.16 |
| *Foxo1* | 1.00 ± 0.05 | 0.96 ± 0.05<sup>†</sup> | 0.57 |
| *Gsk3b* | 1.00 ± 0.04 | 0.96 ± 0.05<sup>†</sup> | 0.58 |

Data are presented as mean ± SEM. *P* values were obtained using an unpaired Student's *t*-test.
<sup>\*</sup> Indicates data not normally distributed and analysed with a non-parametric Mann-Whitney U-test.
<sup>†</sup> *n* = 8.

## Discussion

This study demonstrates for the first time that a low, acute dose of prenatal EtOH can program metabolic dysfunction in a sex-specific manner. Male offspring developed insulin resistance at 6 months of age, regardless of the lack of traditional hallmarks of low birth weight and catch-up growth seen in fetal programming. PAE male offspring also showed alterations in insulin signalling in adipose tissue. Given that BAC levels following alcohol exposure were relatively moderate, at ∼0.05%, these results strengthen the message that no amount of alcohol during pregnancy is safe for long-term offspring health.

Previous preclinical studies investigating the association between PAE and offspring metabolic health have typically used a high dose (2 g/kg BW administered twice a day) throughout gestation and found this can induce glucose intolerance and insulin resistance (see (Akison *et al*., 2019) for review). This dosage typically results in a peak BAC of ∼0.1-0.15% (Chen & Nyomba, 2003b). However, our moderate, acute dose model, resulting in a BAC of just 0.05% one hour after gavage, is more representative of drinking practices in pregnant women who report drinking on average 1-2 standard drinks sporadically during pregnancy. Although the model used in this study has been previously used by Gray et al. (Gray *et al*., 2010), in the same species and strain, they reported a BAC of 0.107% in treated dams 1 h after gavage. The reasons for these differences in BAC are unknown but given these discrepancies, this highlights the importance of measuring and reporting BAC levels, in addition to providing the dosage administered.

In our moderate exposure model, there were no changes in fasting blood glucose levels at any age, or fasting insulin levels in adolescent offspring exposed to prenatal alcohol. Blood glucose concentrations during a GTT performed at 6 months of age were also not altered. However, the major finding of our study was that male offspring exposed to alcohol showed evidence of insulin resistance at 6 months of age, with elevated fasting plasma insulin, resulting in an increased HOMA-IR index, indicative of insulin resistance, and a decreased QUICKI index, indicative of reduced insulin sensitivity.

There was also a significant impairment in glucose clearance following an insulin challenge. These are hallmarks of a pre-diabetic state. The trend for increased first phase insulin secretion during the GTT in PAE males suggests an appropriate compensatory response to this reduced insulin sensitivity, resulting in maintenance of blood glucose to the same level as controls. In a recent systematic review, 12 preclinical studies that examined glucose metabolism using a GTT and/or ITT found evidence of insulin resistance in all but one study (Akison *et al*., 2019). However, the majority of these studies also reported changes in circulating glucose levels following PAE. Interestingly, only one of these 12 studies had a low BAC (peak of ∼0.03%), comparable to that measured in our study. That study, using a chronic low dose of alcohol throughout pregnancy, also found increased first phase insulin secretion in response to a glucose challenge exclusively in males, without concomitant changes in blood glucose levels (Probyn *et al*., 2013). This suggests that the BAC may need to reach a higher threshold to induce changes in glucose tolerance, while changes in insulin sensitivity can potentially manifest following exposure to a much lower BAC. Alternatively, aging may be an important ‘2^nd^ hit’ for a more overt, diabetic phenotype. Adolescent (PN30) and early adult (10-11 week-old) offspring in our study did not have any significant metabolic phenotype, and it was only at 6 months of age when insulin resistance was detected. We predict that examination of offspring further aged to 12 or 18 months may also exhibit glucose intolerance, as well as insulin resistance, although this needs to be tested.

We explored potential molecular mechanisms underlying the observed insulin resistant phenotype. Given the importance of the liver for glucose homeostasis, we examined the expression of a selection of hepatic genes involved in glucose transport (*Glut2*), glucose homeostasis (*G6pc, Gck, Pck1, Ppargc1a*) and insulin signalling (*Insr*). Similar to our low dose chronic alcohol exposure model (Probyn *et al*., 2013), we found no dysregulation of these genes due to PAE, and instead observed only sex-specific differences in gene expression. We also examined AKT protein expression and activation via phosphorylation in the liver but so no evidence of central insulin resistance This suggested the deficits lay within peripheral tissues, skeletal muscle or adipose tissue, with PI3K/AKT signalling known to play an essential role in insulin-mediated glucose uptake in these peripheral tissues (Huang *et al*., 2018). Supporting this, we did find moderate alterations in AKT levels in adipose tissue, both total AKT and activated AKT, measured by phosphorylation at Threonine_308_. Both were significantly elevated in adipose tissue collected from PAE male offspring at 6 months of age. Intriguingly, our results are consistent to a previous study investigating insulin signalling in offspring exposed to periconceptional alcohol (Gardebjer *et al*., 2015). In this previous study, both males and females were insulin resistant but there were inconsistent results with regards to AKT protein levels and activation. In females, AKT levels and phosphorylation were reduced, as you would expect given the phenotype.

However, in EtOH-exposed male offspring, total AKT levels were elevated, as we have seen here. Also, pAKT(Thr_308_) normalised to total AKT was increased in these animals. This suggests that increased total AKT and pAKT(Thr_308_) may be a potential compensatory mechanism, operating exclusively in males, in response to increased fasting insulin levels. This hypothesis requires further investigation. We further interrogated the insulin signalling pathway both upstream (*Insr*) and downstream (*Foxo1* and *Gsk3β*) in the AKT signalling pathway but found no changes in gene expression due to ethanol exposure. We also found no differences in muscle expression of *Glut4* or AKT signalling that have been demonstrated in other models of prenatal alcohol exposure (Chen & Nyomba, 2003a; Chen & Nyomba, 2003b; Chen *et al*., 2005; Yao *et al*., 2006), although this was only seen when tissues were collected immediately following a glucose challenge and not when animals were in a random-fed state. Therefore, our results and others suggest that adipose tissue is the site of insulin resistance in male offspring in response to prenatal alcohol exposure, although further work is needed to clarify the nature of this relationship.

Maternal malnutrition, even if relatively modest, is an established prenatal perturbation which may result in fetal growth restriction and offspring metabolic disorders (Flynn *et al*., 2013; Gao *et al*., 2014). However, this can be excluded in our study, as food and water consumption throughout pregnancy did not differ between the EtOH and control dams. Additionally, there were no differences in other maternal parameters known to affect offspring metabolic outcome, such as obesity (Whitaker, 2004; El-Gilany & Hammad, 2010; Janjua *et al*., 2012), gestational weight gain (Margerison Zilko *et al*., 2010) and blood glucose (used as an indicator for gestational diabetes) (Van Wootten & Turner, 2002). Previous preclinical studies have reported reduced birth weight and subsequent ‘catch-up’ growth in offspring exposed to prenatal alcohol that was associated with subsequent metabolic dysfunction, specifically insulin resistance, in adults (Chen & Nyomba, 2003b; Yao *et al*., 2006; Dobson *et al*., 2014; Xia *et al*., 2014; Gardebjer *et al*., 2015). However, other studies highlight that growth restriction is not required to program chronic disease, with insulin resistance arising in PAE offspring irrespective of birth weight (Chen & Nyomba, 2004; Yao & Nyomba, 2008; Probyn *et al*., 2013; Yao *et al*., 2013). Clinical studies have also reported that growth restriction is not a prerequisite for development of metabolic syndrome (Euser *et al*., 2010; Kopec *et al*., 2017). We also found no effect of PAE on offspring weight or growth, and yet EtOH-exposed males developed insulin resistance at 6 months of age. This highlights that even low-dose exposures can program offspring disease, irrespective of birth weight.

Although our low-dose PAE model did not affect body condition, assessed from morphometric analysis, at 6 months of age, we did not directly measure body fat. Studies have previously assessed body composition in PAE offspring using dual-energy X-ray absorptiometry (DEXA) and/or magnetic resonance imaging (MRI), with variable results. Our result is similar to that found in a previous low-dose model in rats (Probyn *et al*., 2013), as well as a study in mice with EtOH exposure restricted to late gestation (Amos-Kroohs *et al*., 2018). However, other higher dose models have reported increased adiposity in PAE offspring compared to controls, particularly in males (Dobson *et al*., 2012; Gardebjer *et al*., 2018; Zhang *et al*., 2018).

PAE offspring at 4 months of age did not exhibit a preference for a HFD. However, they did eat slightly less than controls, which resulted in a significantly lower energy consumption in PAE offspring due to the HFD being more energy rich. This was particularly evident in the second choice period, when the novelty of the new diet had presumably waned (Dorey *et al*., 2018). This was contrary to a recent study in rat offspring exposed to a higher dose of alcohol exclusively around conception, which found that males preferentially consumed more HFD than controls (Dorey *et al*., 2018). However, this study was conducted in much older rats (15 months of age). Clinical studies have found that children diagnosed with FASD tend to have hyperphagia and abnormal eating patterns, including constant snacking and impaired satiety (Werts *et al*., 2014; Amos-Kroohs *et al*., 2016). Our results are more consistent with a study conducted in mice, which found no changes in food preference as a result of PAE (Amos-Kroohs *et al*., 2018). Interestingly, that study also restricted PAE to later in gestation (E12.5-17.5). Taken together, these studies and ours suggest that a change in eating behaviour is not consistently associated with prenatal alcohol exposure but may relate to the timing and severity of the exposure as well as offspring age. Our study also suggests that altered food consumption or a propensity for consuming a HFD does not underlie the metabolic dysfunction observed in male offspring. Future studies examining the impact of PAE on offspring metabolism should include analysis of feeding behaviours to further define any associations.

A possible limitation of this study was that the saline gavage did not control for the additional energy provided by the EtOH. A recent study highlighted that the typical maltodextrin control, used as a substitute for the calories provided by EtOH, is not metabolised in the same way, and when a medium chain triglycerides solution was included as a control, no changes in metabolic outcomes could be attributed to PAE (Amos-Kroohs *et al*., 2018). However, our dose of EtOH per gavage was so small (∼9-10 kJ), that when we calculated the combined energy intake attributed to the EtOH and chow on the days of gavage, there was no significant difference between the EtOH and Control dams in our study. Therefore, our saline treatment adequately controlled for the potential effects of the gavage treatment and did not result in a lower overall energy intake when compared with our low-dose EtOH treatment group.

## Conclusions

This study supports the programming of offspring metabolic health as a result of a relatively moderate PAE during a critical period of development. Although ethanol was administered during mid-pregnancy in the rat, this equates to ∼week 10-12 of a human pregnancy when a woman may not yet be aware she is pregnant, particularly if the pregnancy was not planned. The pre-diabetic, insulin resistant phenotype reported in male offspring of PAE dams occurred despite a relatively low BAC and standard postnatal care, and without the ‘2^nd^ Hit’ of a high fat diet as used by others to ‘unmask’ disease. Programmed alterations were also independent of maternal parameters known to affect offspring metabolic health, such as gestational diabetes and obesity. Therefore, this study highlights the importance of abstaining from alcohol consumption during pregnancy or when planning a pregnancy.

## Additional information

### Author contributions

All experiments were performed in the Developmental Programming in Disease Lab, School of Biomedical Sciences, The University of Queensland. K.M.M. and L.K.A. were responsible for the conception and design of the experiments. T.M.T.N., L.K.A. and S.E.S. were responsible for the collection of the data. All authors were involved in data analysis and/or interpretation. All authors contributed to drafting the work and/or revising it critically for important intellectual content. All authors approve of the final version of the manuscript and agree to be accountable for all aspects of the work in ensuring that questions related to the accuracy or integrity of any part of the work are appropriately investigated and resolved. All persons designated as authors qualify for authorship, and all those who qualify for authorship are listed.

## Acknowledgements

We would like to acknowledge Natasha Steiger (Animal Endocrinology Lab, School of Biomedical Sciences, University of Queensland) for analysis of plasma insulin; Dave Herne and Barb Arnts (University of Queensland Biological Resources) for assistance with animal treatments and husbandry; Elizabeth McReight (School of Biomedical Sciences, University of Queensland) for assistance with animal work; and Jacobus Ungerer (Pathology Queensland, Queensland Health) for analysis of BAC.

## Supporting information

All underlying data used in figures and tables reported in this study and can be accessed from the following URL: https://doi.org/10.5281/zenodo.3406863.

